# Sex matters: effects of sex and mating in the presence and absence of a protective microbe

**DOI:** 10.1101/2021.05.24.445428

**Authors:** Anke Kloock, Lena Peters, Charlotte Rafaluk-Mohr

## Abstract

In most animals, female investment in offspring production is greater than for males. Lifetime reproductive success (LRS) is predicted to be optimised in females through extended lifespans to maximise reproductive events. Extended lifespan can be achieved through increased investment in immunity. Males, however, maximise lifetime reproductive success by obtaining as many matings as possible. Microbe-mediated protection (MMP) is known to affect both immunity and reproduction, but whether the two sexes respond differently to the provision of MMP remains to be explored. Here, we investigated the sex-specific differences in host life history traits between female and male *Caenorhabditis elegans* following pathogenic infection with *Staphylococcus aureus* with or without MMP by *Enterococcus faecalis*. Overall, female survival decreased with increased mating. With MMP, females increased investment into offspring production, while males displayed higher behavioural activity. These results highlight the different strategies employed by the two sexes under pathogen infection with and without MMP.

## Introduction

Mating is costly. The costs paid by females and males, however, differ (Jens Rolff, 2002; Jens Rolff et al., 2005). Females experience higher costs for the provision of gametes, as eggs are highly costly (Stearns, 1987) and risk potential harm during mating (Le Page et al., 2017; Rankin & Kokko, 2006). Males on the other hand, generally must invest more into outcompeting other males by male-male competition (Le Boeuf, 1974) or searching for mates (Lipton et al., 2004). These traits are energetically costly, and as such it is hypothesized that there are trade-offs between survival and reproduction in both females and males (Sheldon & Verhulst, 1996), albeit differentially mediated in the two sexes.

The need for females to spread reproduction across their lifetime is expected to result in differential investment in immunity, an effect that has been demonstrated across taxa. Following infection males of *Panorpa vulgaris, Caenorhabditis elegans* and *Daphnia magna* have higher infection load, reduced lysozyme-like activity and haemocytes, which results in reduced survival and body size (Kurtz et al., 2000; Masri et al., 2013; Roth et al., 2008). *C. elegans* males do, however, show higher activity (observed as higher escape behaviour) (Masri et al., 2013), which is linked to behavioural avoidance (Schulenburg & Ewbank, 2007). In contrast, female scorpion flies have increased immune activity (Kurtz et al., 2000) and women survive pandemics (Zuk, 2009), slavery or famines better than men do (Zarulli et al., 2018).

Given these differences, we hypothesized that the two sexes would benefit differently from a protective microbe. Protective microbes can be important in host defence in the face of infection, a phenomenon referred to as “defensive mutualism” (King, 2019), where microbes can supplement the host’s immune system (Abt & Artis, 2013). Defensive mutualisms have been observed across kingdoms (reviewed in (Ford & King, 2016)). The potential of defensive mutualism to enhance survival as well as offspring production has been observed repeatedly (King et al., 2016; Kloock et al., 2020; Koehler et al., 2013). However, most of these examples have only considered population-level effects, while few studies have focussed on individual behaviours and/or sex differences between the hosts (McLean et al., 2018).

To test for differences in life history traits between the two host sexes with or without microbe-mediated protection (MMP) during infection, we used an established experimental system using *Caenorhabditis elegans* as a host, *Enterococcus faecalis* as a protective microbe and pathogenic *Staphylococcus aureus* (Ford et al., 2016; King et al., 2016; Rafaluk-Mohr et al., 2018). Here, we used a population of *C. elegans* made up of males and feminised hermaphrodites, that only carry eggs and cannot produce sperm, and thus are referred to as females (Theologidis et al., 2014). *C. elegans* males are known to actively search for females to mate with and in the absence of females this behaviour is increased (Lipton et al., 2004). For our experiment, we separated females and males and manipulated the time frame that they could come into contact with one another and mate. Individuals were either unmated, short-term mated or lifetime mated. To test the impact of MMP, worms were offered one of three bacterial diets: food only, pathogen or pathogen with MMP. We investigated the differences in life history traits for the two sexes under the different mating and bacterial diet conditions.

## Materials and Methods

### Worm and bacteria system

We used an obligate outcrossing worm population (line EEVD00 from Henrique Teotonio (Theologidis et al., 2014)) where worms carry the *fog-2(q71)* mutation, preventing hermaphrodites from producing sperm (Theologidis et al., 2014). Worms were kept on Nematode Growth Medium (NGM) (Brenner, 1974), and fed non-pathogenic *Salmonella*, hereafter referred to as food (Desai et al., 2019; Diaz et al., 2015; Kloock et al., 2020). For pathogenic infection, the Gram positive *Staphylococcus aureus* strain MSSA476 (Holden et al., 2004) was used. The strain OG1RF of *Enterococcus faecalis* (Garsin et al., 2001) was used as a protective microbe against *S. aureus* infection (Ford et al., 2016; King et al., 2016; Rafaluk-Mohr et al., 2018).

### Pathogenic infection and long-term survival analysis

All assays were carried out blind. Worms were sterilised and synchronized via bleaching (Stiernagle, 2006). Simultaneously, the bacteria were grown in overnight cultures: Either *E. faecalis* overnight in 25ml of Todd-Hewitt Broth (THB), or food in 25ml of Lysogeny broth (LB), both at 30°C in a shaking incubator. 6cm NGM plates were inoculated with either 400µl of food or 200µl of food mixed with 200µl of *E. faecalis*. 600 L1 worms were added to each NGM plate and kept at 20°C for 42h. Simultaneously, a liquid culture of *S. aureus* was grown in THB from a frozen stock, while food was grown in LB. Both cultures were incubated under shaking conditions at 30°C overnight. The following day, 20µl of *S. aureus* overnight culture was pipetted onto 3cm on Tryptone Soy Broth agar (TSB) plates and incubated at 30°C overnight. Simultaneously, 6cm NGM plates were inoculated with 150µl food. These plates were used to split worms into groups of only females, only males or 50:50 mixed for 6-8 h (time point when the first eggs appeared on the plate) (Table S1). After worms had mated, 50 worms were placed onto the *S. aureus* lawn with a platinum wire pick and left at 25°C for 24h (Table S1) in groups of unmated, short-term mated and lifetime mated individuals of both sexes.

Survival upon pathogenic infection was scored after 24h. Worms were considered dead if they did not respond to touch with a platinum wire pick. After survival was scored, 10 worms were transferred to 3cm NGM plates seeded with 150µl food and placed at 25°C. Worms were transferred to new plates every 24h with a platinum wire until no further offspring production occurred. Survival was scored every day until all worms were dead. For the food alone treatment, the long-term survival assay followed a similar protocol except that the experimental procedure was carried out at 20 °C, as is standard for *C. elegans* (Amrit et al., 2014).

### Activity analysis

After 24h on the pathogen infection plates, worm behavioural activity was determined via calculating the fraction of worms at the edge of the plate. Worms were considered at the edge of the plate, if they could not be seen from above.

### Avoidance analysis

The proportion of missing worms was calculated 24h after pathogen infection and at each transfer in the long-term survival analysis, as previously described (Pees et al., 2017). The last time point of each experiment was plotted.

### Offspring production

The presence or absence of viable offspring on a plate was noted during pathogen infection and during long term survival. Offspring production was defined as a proportion of plates that had viable offspring over the total amount of plates per treatment.

### Statistical Analysis

Statistical analyses were carried out with RStudio (Version 1.1.463 for Mac)(RStudio Team, 2020). Figures were created with the ggplot2 package (Version 2.1.0). All data, except for the long-term survival and offspring production data, were analysed with nested binomial mixed effects models (GLMM - R package lme4) (Bates et al., 2015, p. 4) to test for an effect of sex, the mating status or an interaction between the two. If the interaction had a significant effect, a Tukey multiple-comparison tests (R package multcomp) (Hothorn et al., 2008) was performed. The long-term survival data was analysed with Kaplan Meier Log Rank test with FDR correction for multiple testing (Therneau, 2020; Therneau & Grambsch, 2002). The offspring data was analysed using a Wilcoxon Rank Test (Mann & Whitney, D. R., 1947).

## Results & Discussion

### Lifetime mated females are most affected by pathogen infection and mating

During pathogen infection without (Figure 1A) and with MMP (Figure 1B), males survived better than females (p<0.001, GLMM, X^2^=121.171, df=1 and p<0.001, GLMM, X^2^=20.864, df=1, respectively), while over a lifetime this pattern was only present without MMP (Figure 1C-E; without MMP: p<0.001, Kaplan Meier Survival Estimates (KMSE); on food: p=0.62 (KMSE), with MMP: p=0.32, (KMSE)). Females were dramatically affected by the mating status, during pathogen infection with and without MMP (p<0.001, GLMM, X^2^=84.252, df=2 (Figure 1A) and p<0.01, GLMM, X^2^=10.759, df=2 (Figure 1B), respectively) and also over a lifetime independent of the bacterial diet (Figure 1C-E), where lifetime mated females survived worse than their unmated or short-term mated counterparts (on food: both p<0.001, (KMSE); without MMP: both p<0.001 (KMSE); with MMP both p<0.001 (KMSE)). Males were not affected by mating, neither during pathogen infection nor over a lifetime (all p>0.05).

**Figure 1:**
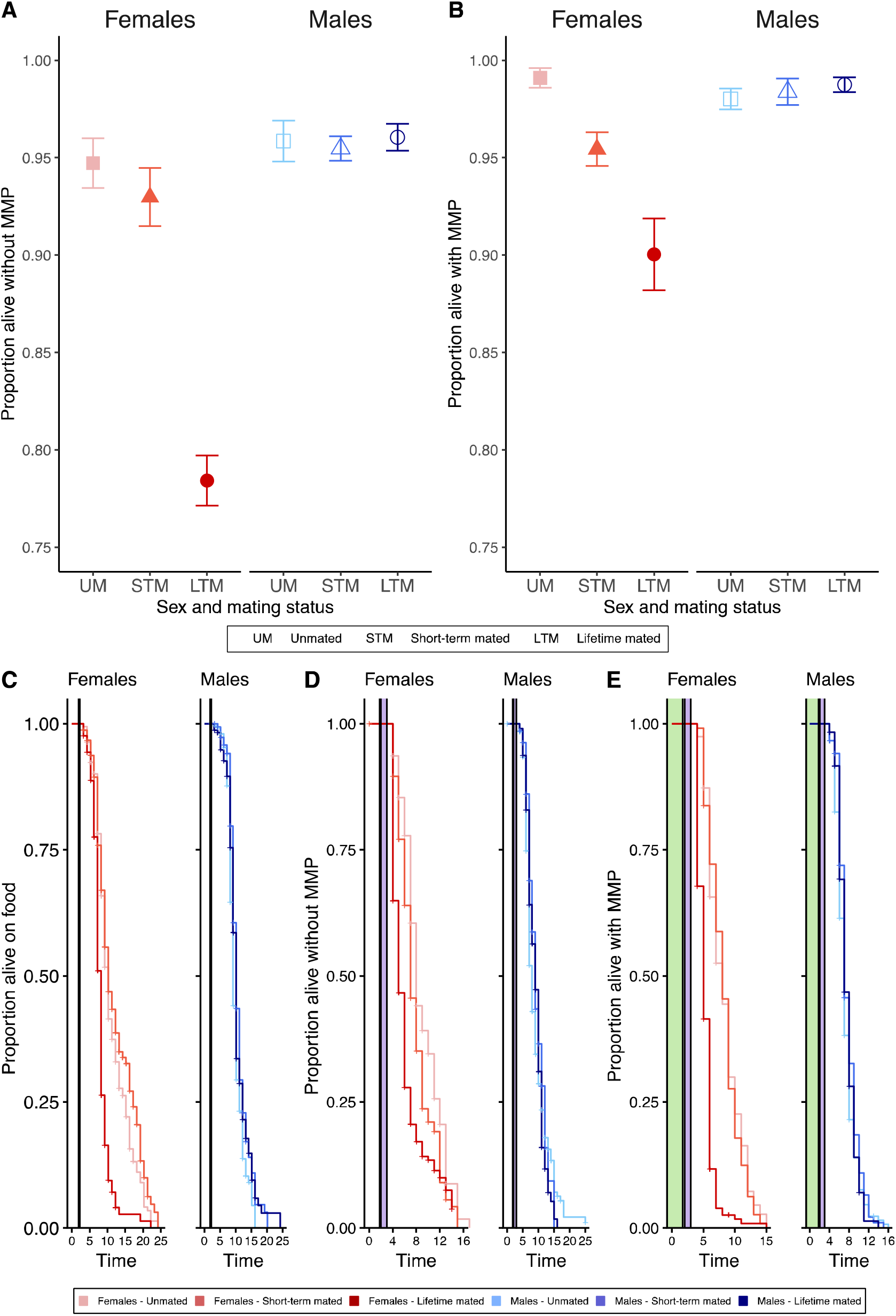
Survival of different sexes and mating treatments after 24h pathogen infection (A, B) and lifelong (C-E). (A) Without MMP females suffer more from mating than males do, while males survive better overall. (B) With MMP, females are suffering more from mating, while males survive better overall. (C) When only ever being exposed to food (in white), lifetime mated females survive worse than any other females, while only lifetime mated males survive worse than short-term mated males. (B) After pathogen infection (in purple) without MMP, lifetime mated females survive worse than short-term mated females, which survive worse than unmated females. Male survival was not affected by mating, while males live overall longer than females do. (C) After pathogen infection with MMP (in green), lifetime mated females survive worse than short-term mated females, while males are not affected by mating. (A, B) Each point represents the mean ± the standard error of the mean of four biological replicates and three or four technical replicates. (C-E) Each curve represents the Kaplan Meier Survival estimate for three or four technical replicates and four or five biological replicates. Vertical lines indicate when worms were transferred to a new bacterial diet.

Females suffer worse the longer they have been mated with males, while males survive pathogen infection better than females independent of MMP (Figure 1A & 1B). This pattern holds during pathogen infection (Figure 1A & 1B) and over a lifetime (Figure 1C-E). The potential of MMP to enhance survival (King et al., 2016; Kloock et al., 2020; Martinez et al., 2016) as well as offspring production (Koehler et al., 2013) has been shown repeatedly. So far, these effects have mainly been considered at the population level. However, the role of MMP might have different effects on individual behaviours of different sexes in different life stages (McLean et al., 2018).

A potential explanation for the observed phenotype could be mechanical gut integrity, which can be different between males and females as observed in *Drosophila* (Regan et al., 2016). The pathogen used here, *S. aureus*, is known to accumulate in the worms gut and to kill worms by distention of the intestinal lumen (Sifri et al., 2003). If gut integrity would thus be more easily damaged in *C. elegans* females, but not in males (personal observation, Figure S2), this could serve as a potential explanation as to why females are more harshly affected by pathogenic infection with *S. aureus*. The potentially disrupted gut integrity by *S. aureus* infection, could further be weakened by mechanical penetration by males, which would explain why lifetime mated females survive worse during pathogen infection and over a lifetime.

The act of mating is costly and life shorting (Gems & Riddle, 1996), independent of offspring production, as here short-term mated females, that also produce costly offspring, do not have lower survival than unmated females over a lifetime (Figure 1C-E). Although previous studies in *C. elegans* have not demonstrated a cost for the timespan of mating (Booth et al., 2019), these studies have limited mating to 12 hours (Booth et al., 2019; Shi et al., 2019), while our worms are only parted by death. We saw the greatest costs when worms had the opportunity to mate for their entire lifespan suggesting that differences could have been masked by restricted time windows for mating in previous studies. Furthermore, we used an obligately sexual strain (Theologidis et al., 2014), where females had to mate to reproduce, which could have potentially resulted in increased willingness to mate in females, and also led to the lack of self-sperm, which protects self-fertilising hermaphrodites from male induced demise (Booth et al., 2019; Shi et al., 2019; Shi & Murphy, 2014). During mating males transfer seminal fluids alongside sperm, which reduces female life span in mice, *Drosophila*, and *C. elegans* (Booth et al., 2019; Chapman & Partridge, 1996; Partridge & Gems, 2002), and can leave females immuno-supressed post mating (J. Rolff & Siva-Jothy, 2003).. Our results reflect a wealth of findings in other species that has shown mating to be costly, such as in Drosophila (Fowler & Partridge, 1989), or birds (Liker & Székely, 2005).

### Male and female behaviour is linked to the presence of the other sex

Females and males respond differently to infection and thus also display different behaviour in the face of infection. Those plates with both sexes present during pathogen infection were more active, measured here as the proportion of worms at the edge (Figure 2A), independent of MMP ((p<0.001, GLMM, X^2^=122.56, df=5 and p<0.001, GLMM, X^2^=51.8544, df=2, respectively; Figure 2 B, C). As this increased activity for lifetime mating worms can be a hint to increased avoidance behaviour, which is itself a mechanism to respond to pathogenic infection (Pees et al., 2017), we also assessed the proportion of missing worms during pathogen infection (Figure D, E) and over a lifetime (Figure 2F-H). Males went missing with a higher proportion than females during pathogen infection independent of MMP (without MMP: p<0.001, GLMM, X^2^=17.5919, df=1 and with MMP: p<0.001, GLMM, X^2^=35.586, df=1) and over a lifetime on food alone (p<0.001, GLMM, X^2^=154.21, df=1) and with MMP (p=0.002, GLMM, X^2^=9.198, df=1), while the difference between the sexes was not significant after pathogen infection without MMP (p=0.23, GLMM, X^2^=1.42, df=1). Interestingly on food alone, lifetime mated worms displayed opposite patterns in comparison to unmated worms: for females there were more lifetime mated females than unmated females missing (p<0.01, GLMM, X^2^=201.31, df=5), while for males there were less lifetime mated males than unmated males missing (p<0.01, GLMM, X^2^=201.31, df=5).

**Figure 2:**
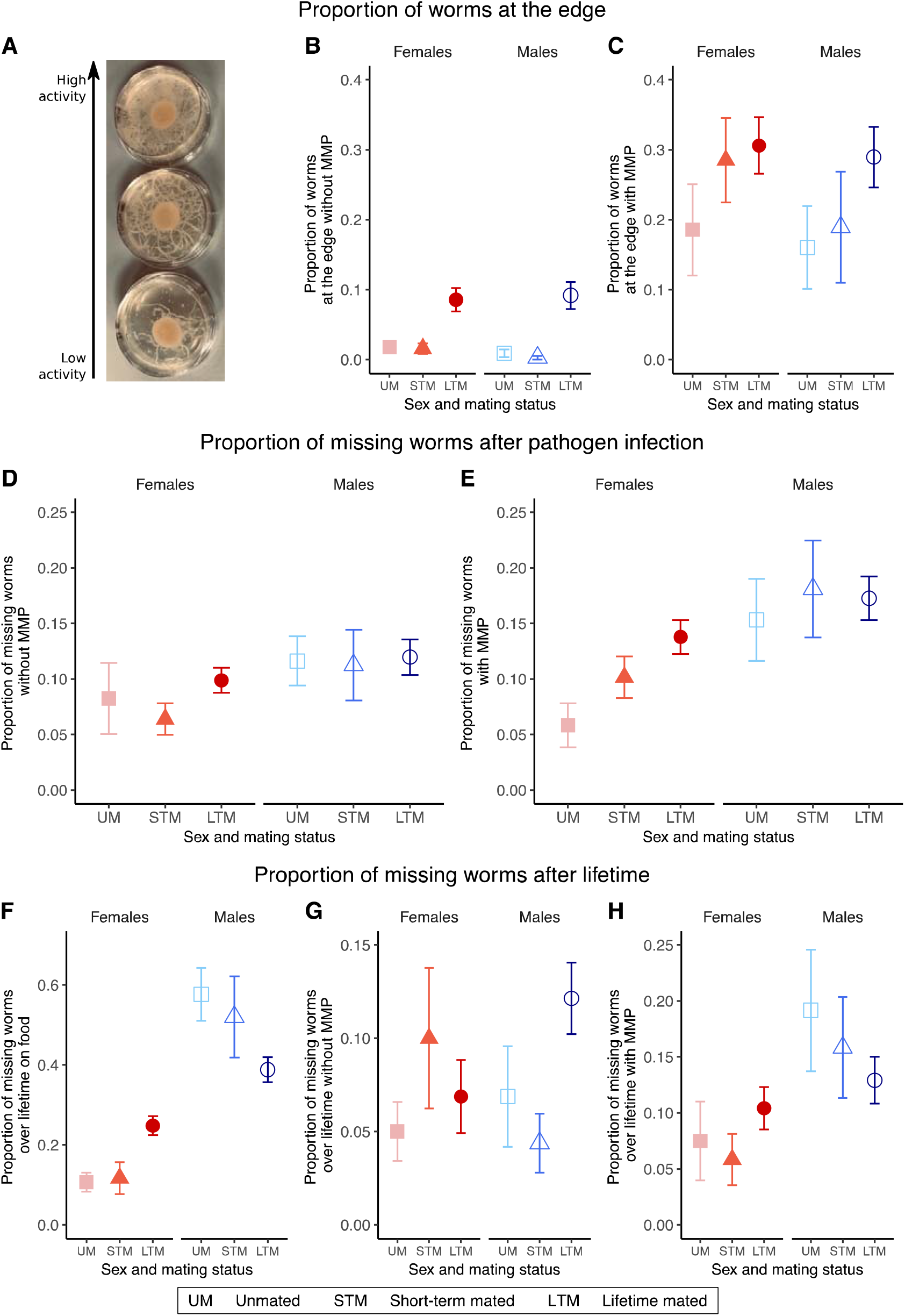
Female and male behaviour is linked to the presence of the other sex. (A) Picture of three pathogenic plates with different activity levels from low to high. (B) Without MMP after pathogen infection plates with lifetime mated worms of both sexes had a higher proportion of worms at the edge of the plate. (C) With MMP after pathogen infection, plates with lifetime mated worms of both sexes have a higher proportion of worms at the edge of the plate. Proportion of missing worms of different sexes and mating treatments either during pathogen infection (D, E) and over lifetime (F-H, only the last time point is plotted). (D) During pathogen infection without MMP, a higher proportion of males is missing than of females. (E) During pathogen infection with MMP, more males went missing than females. (F) Proportion of missing worms on food over lifetime. Females showed lower proportion of missing worms than males. (G) Proportion of missing worms without MMP over lifetime with no differences detected. (H) Proportion of missing worms with MMP over lifetime, where males have a higher proportion of missing worms than females do. (B-H) Each point represents the mean ± the standard error of the mean of four or five biological replicates and three or four technical replicates. (F-H) Changes in proportion of missing worms over time can be found in Figure S1

Activity level was mainly determined by the mating status, while the proportion of missing worms was predominantly determined by sex. Infected females might thus not be able to move around as much as healthy females, as only healthy females and males on food alone show effects of mating. The mating status also affected the proportion of missing worms in opposite directions in lifetime mated worms on food: females would be missing with a higher proportion with males present while males would stay with a higher proportion with females present. A parsimonious explanation is that males exhibit higher mate searching behaviour and this is increased when females are not present (Lipton et al., 2004). The proportion of missing worms observed here does not appear to be a consequence of pathogen avoidance behaviour (Pees et al., 2017), as this behaviour is not observed for pathogen infection without MMP.

### MMP enables females to invest in offspring production during pathogenic infection

Pre-mated females and non-pre-mated females produced different numbers of eggs. Without MMP, pre-mated females produced more offspring (Wilcoxon Rank Test (WRT), W=57.5, p=0.004, Figure 5A) but not with MMP (WRT, W=18, p=1, Figure 5B). This pattern can however be rescued over a lifetime (Figure S3). It is possible that during pathogen infection females invest great parts of their energy into fighting the infection or are suffering too much from pathogenic infection, and thus invest less in reproduction. This is in contrast to previous findings that show fecundity compensation in response to *S. aureus* infection in *C. elegans* (Pike et al., 2019). This is likely because we only scored for the presence/absence of offspring at a set time point and thus would not be able to detect any change of timing in offspring production, as was described for fecundity compensation in this system of *C. elegans* and *S. aureus* (Pike et al., 2019).

**Figure 5:**
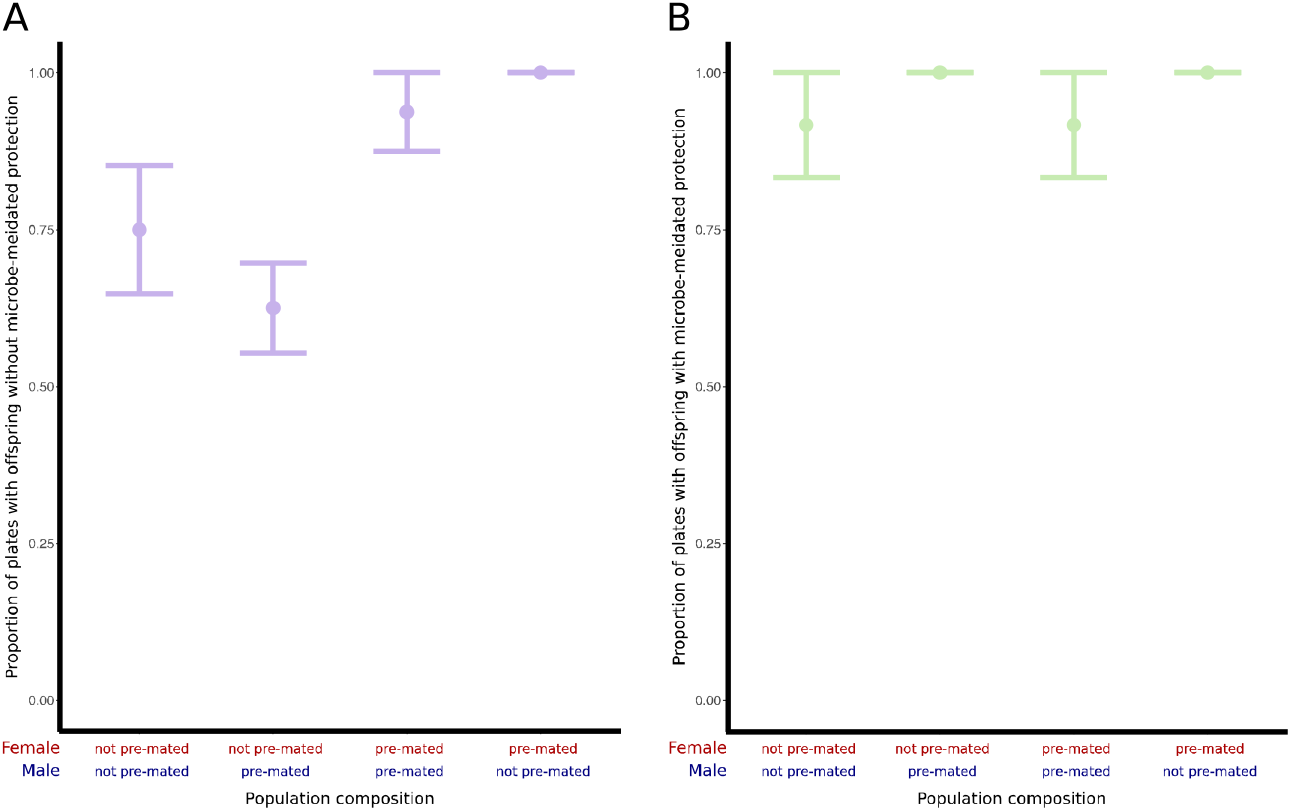
Differences in offspring production during pathogen infection without (A) and with (B) MMP (A). During pathogen infection but without MMP, there are less plates with offspring, despite the presence of males. (B) During pathogen infection, but with MMP, there are no differences between pre-mated and not pre-mated females.

## Conclusion

In conclusion, female survival decreases with increasing mating with males, while male survival was unaffected by mating. This pattern holds over a lifetime and on different bacterial diets. The two sexes benefit from MMP differently. With MMP females invest more energy into egg production, while males invest more into mate searching behaviour. This study highlights that in spite of consistent population level responses to defensive mutualists, individual variation depends heavily on diet, sex, mating status and interaction of these factors. Even though defensive mutualists provide benefits to the host, for females, mating comes at a high cost.

## Acknowledgments

We would like to thank Kayla King for providing funding, laboratory space and intellectual support for the project and the King group for support throughout the laboratory experiments, particularly Maria Ordovas-Montanes and Andrea Gomez Charmorro. We would also like to thank Michael Bonsall for valuable discussion and feedback on the analysis. We are also grateful for reviewer comments.

## Authorship Contributions

AK and CRM conceived and designed the project. AK and LP performed all experiments. AK and CRM performed the statistical analysis. AK and CRM wrote the manuscript and all authors agreed on the final version.

## Supplemental Material

**Supplemental Figure S1:**
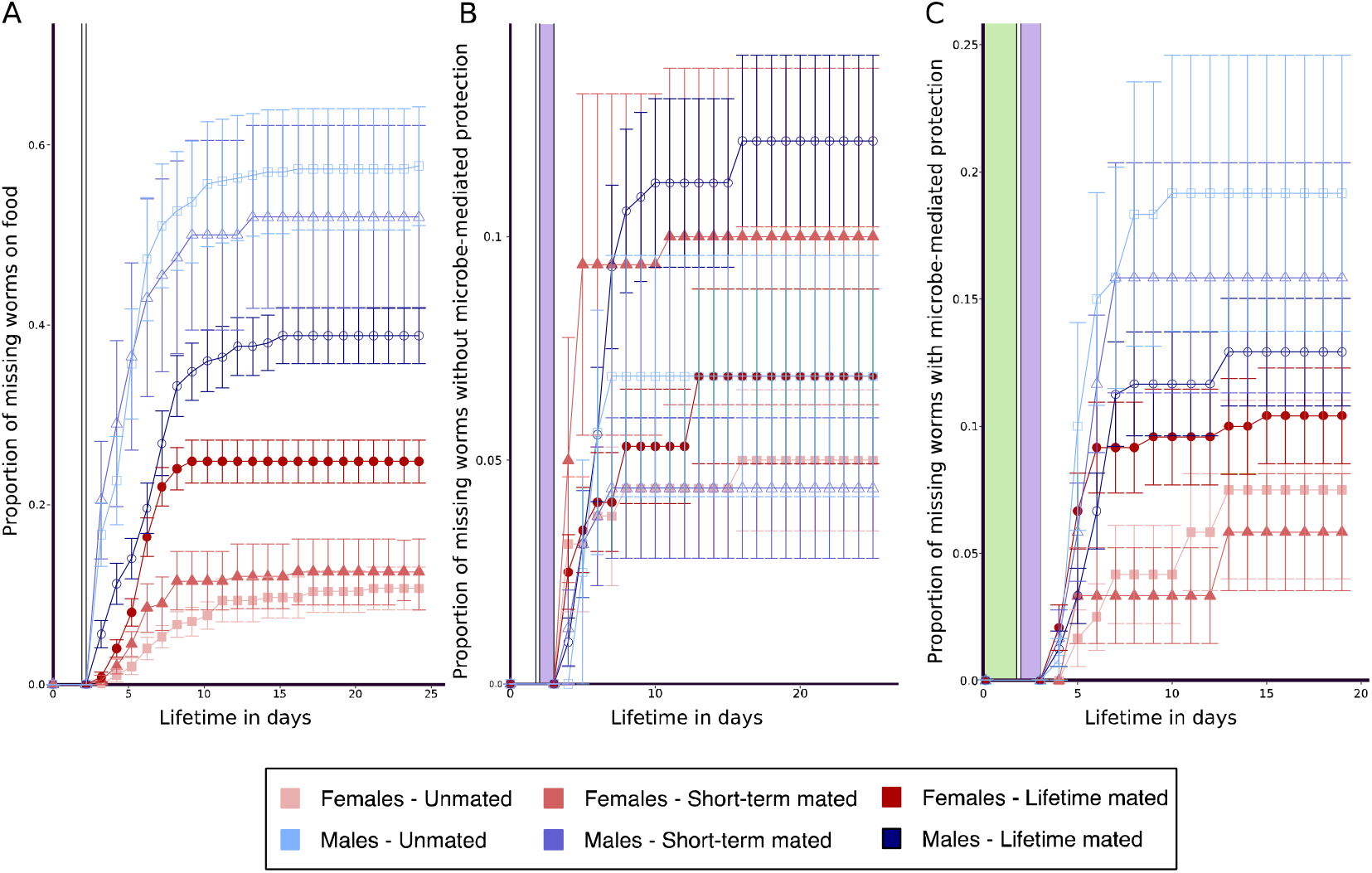
Livelong proportion of missing worms on food (A), with microbe-mediated protection(B) and without of microbe-mediated protection (C). All graphs show the mean +-the standard error of the mean across time.

**Supplemental Figure S2:**
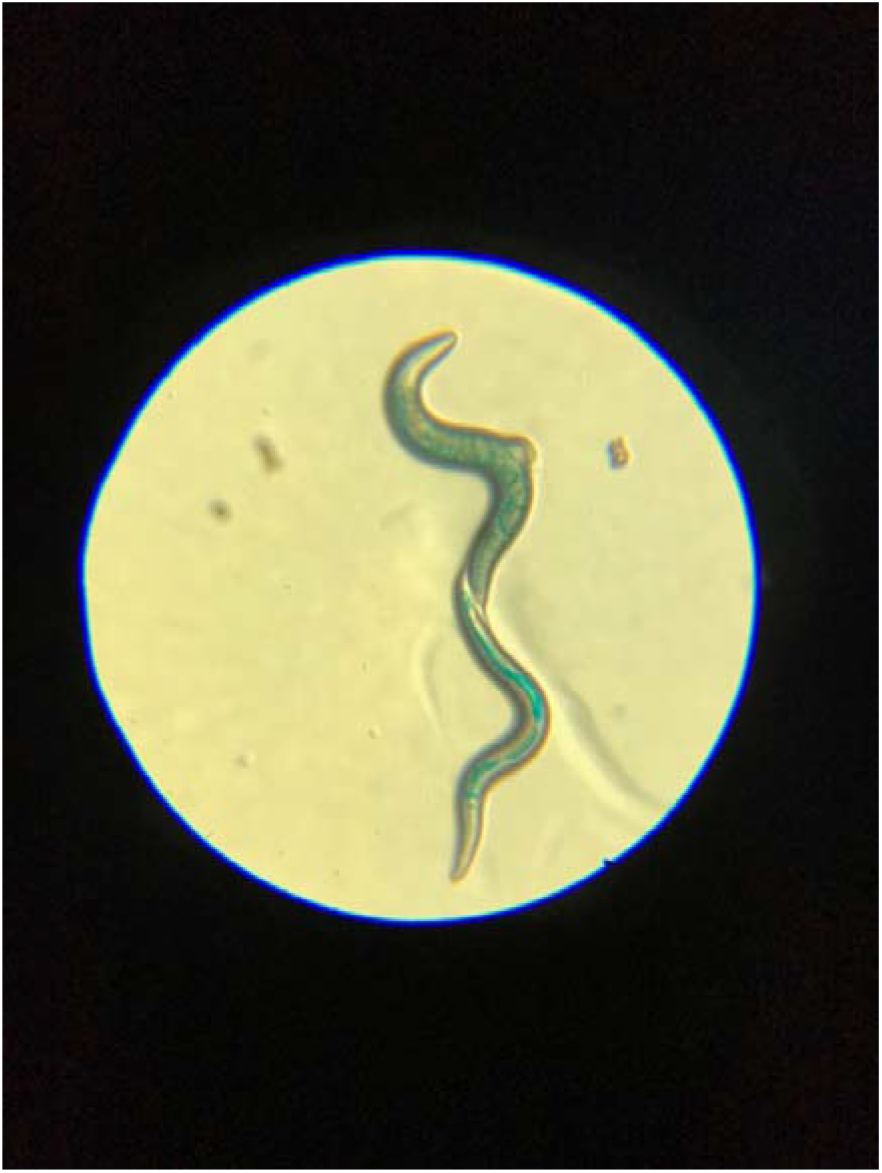
Females show hints of lost gut integrity, while male gut integrity is still intact. Worms that were fed food, coloured with a blue food dye. If gut integrity is still fully intact, blue dye can only be seen in the intestine (as in the lower worm – a male), while if the gut integrity is out of balance, the blue dye can be found in the whole worm body cavity (as in the upper worm – a female).

**Supplemental Figure S3:**
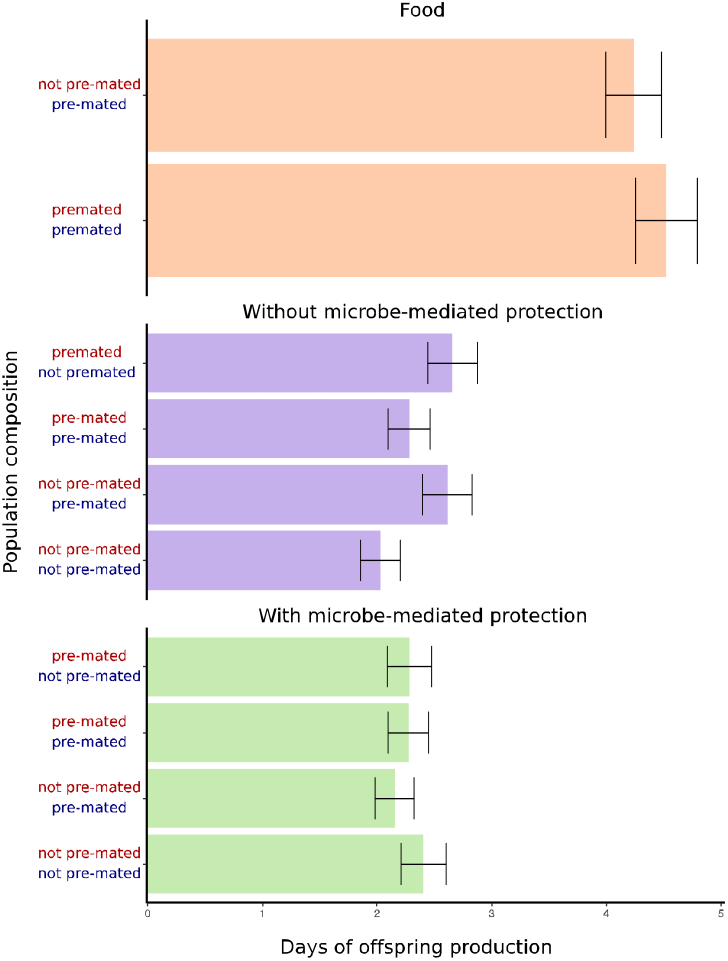
Over a lifetime no difference between pre-mated and not pre-mated females for the days of offspring production can be observed independent on whether worms were raised on food, infected with the pathogen with or without MMP. (A, B) Each point represents the mean ± the standard error of the mean of three or four technical replicates.

**Table S1:**
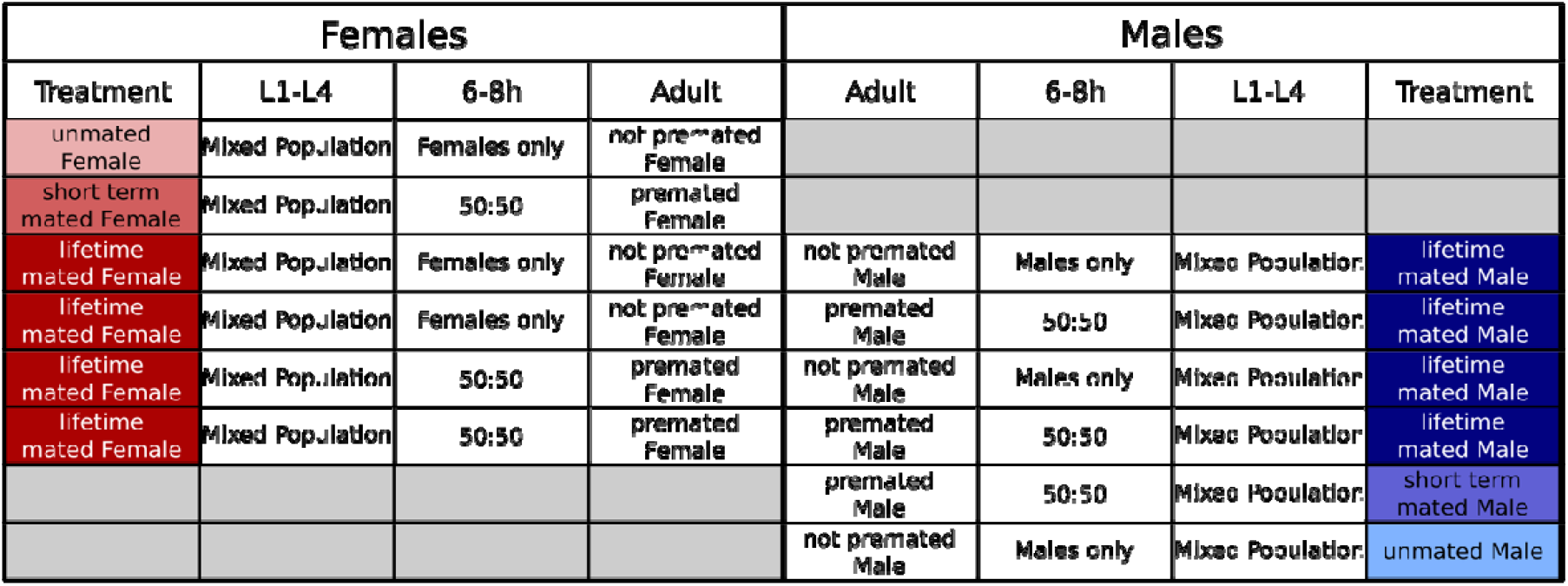
Mating Treatments: Three different mating treatments (Unmated, Short-term mated and Lifetime mated) were set up for both sexes (Female in Red and Male in Blue), which were either single sex plates for each sex (Females only or Males only) or a 50:50 mixed population. Worms were left on these mating plates for 6-8h, before three different mating treatments were set up (unmated, short-term mated, and lifetime mated, the darker the colour the longer the mating period) for each sex.

| Females |  |  |  | Males |  |  |  |
| --- | --- | --- | --- | --- | --- | --- | --- |
| Treatment | L1-L4 | 6-8h | Adult | Adult | 6-8h | L1-L4 | Treatment |
| unmated Female | Mixed Population | Females only | not pre-mated Female |  |  |  |  |
| short term mated Female | Mixed Population | 50:50 | pre-mated Female |  |  |  |  |
| lifetime mated Female | Mixed Population | Females only | not pre-mated Female | not pre-mated Male | Males only | Mixed Population | lifetime mated Male |
| lifetime mated Female | Mixed Population | Females only | not pre-mated Female | pre-mated Male | 50:50 | Mixed Population | lifetime mated Male |
| lifetime mated Female | Mixed Population | 50:50 | pre-mated Female | not pre-mated Male | Males only | Mixed Population | lifetime mated Male |
| lifetime mated Female | Mixed Population | 50:50 | pre-mated Female | pre-mated Male | 50:50 | Mixed Population | lifetime mated Male |
|  |  |  |  | pre-mated Male | 50:50 | Mixed Population | short term mated Male |
|  |  |  |  | not pre-mated Male | Males only | Mixed Population | unmated Male |

**Table S2:**
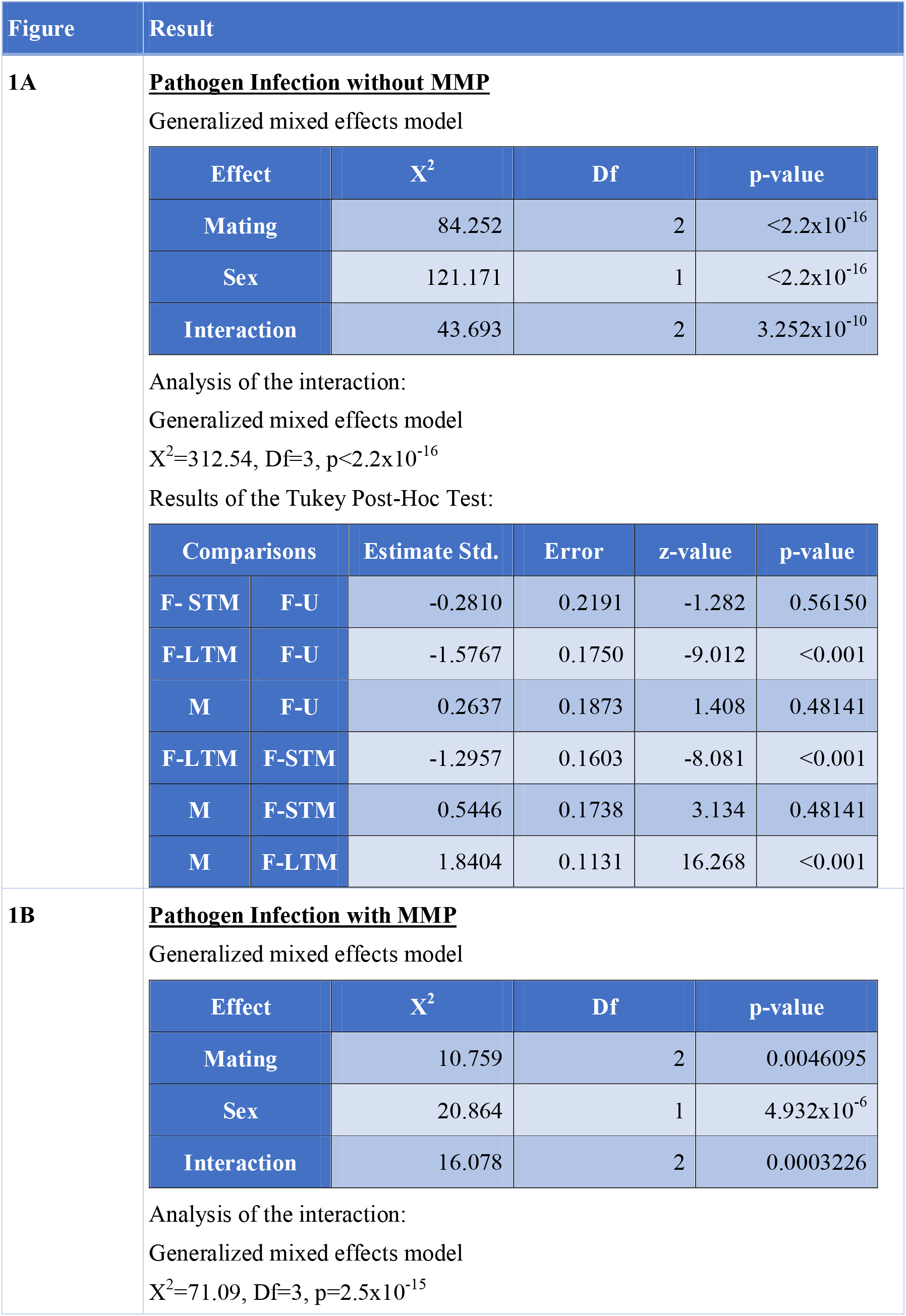
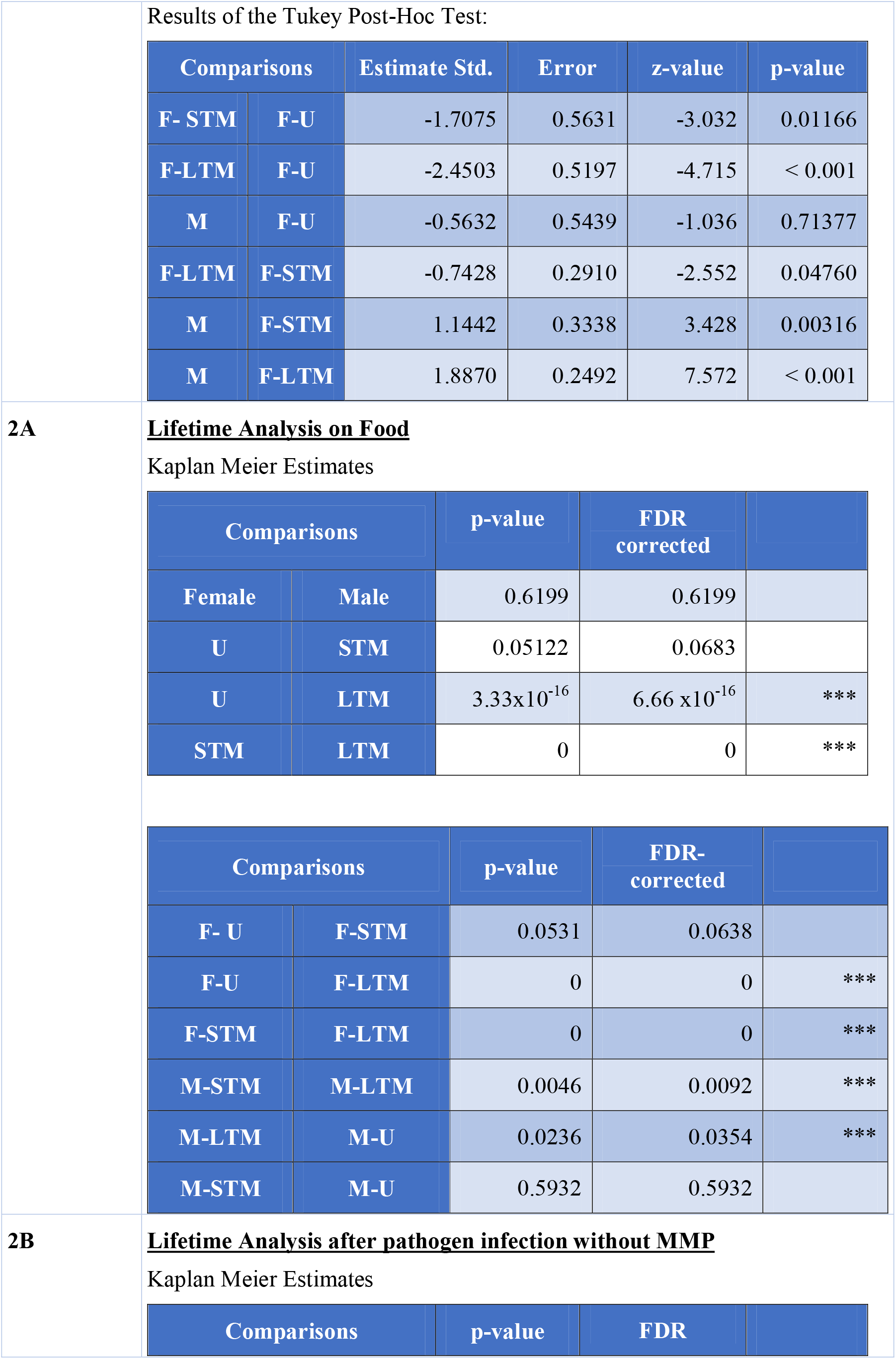
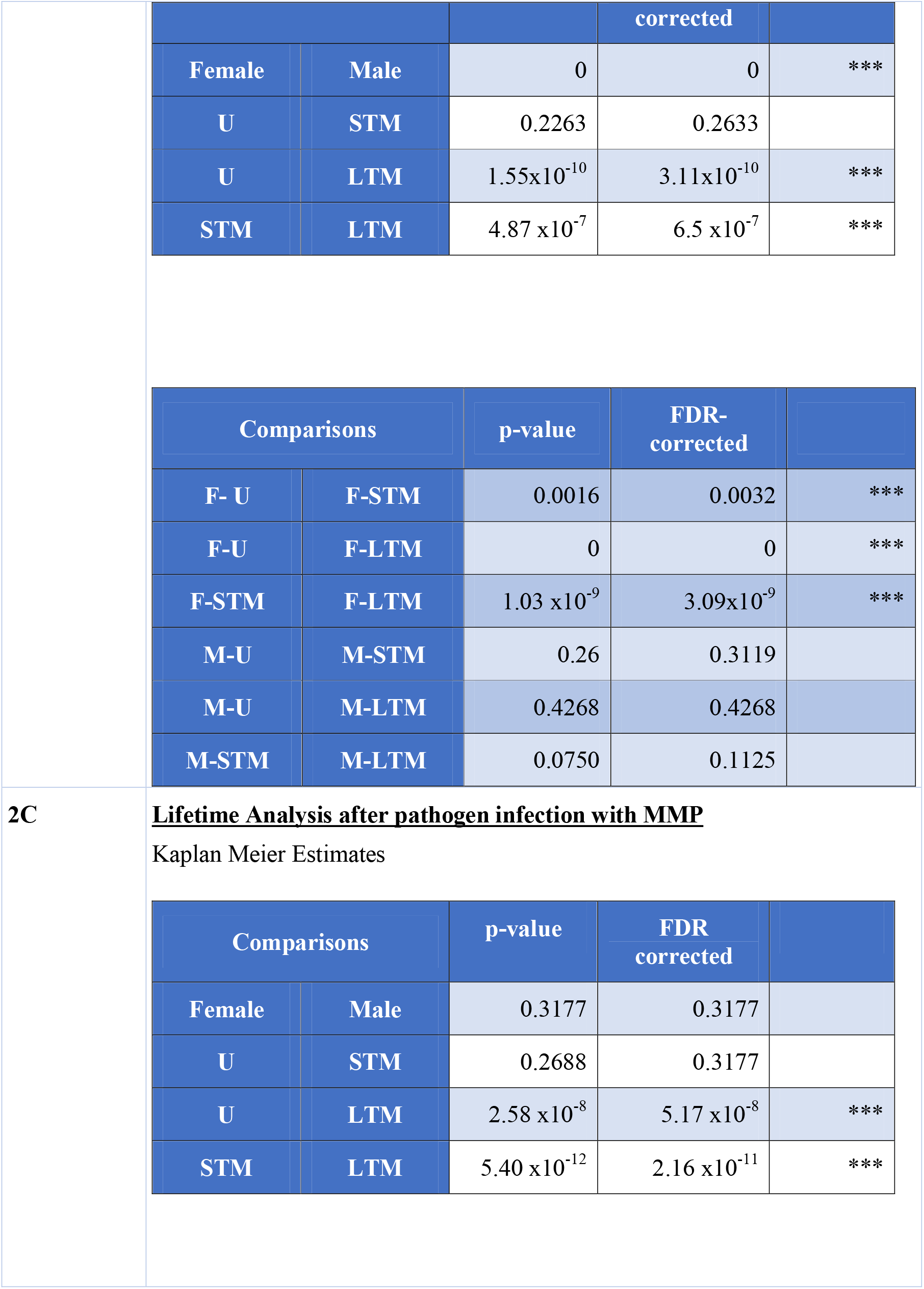
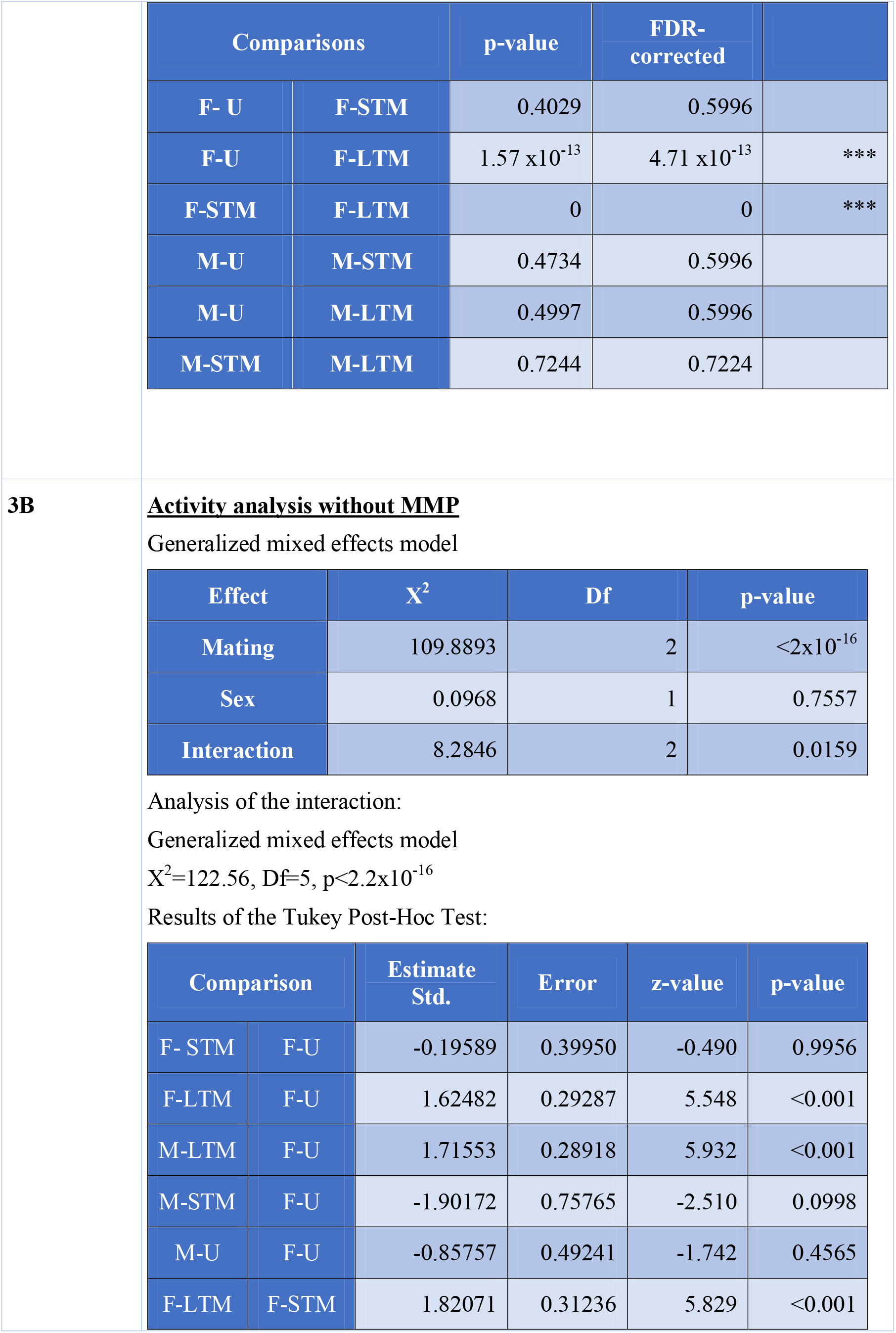
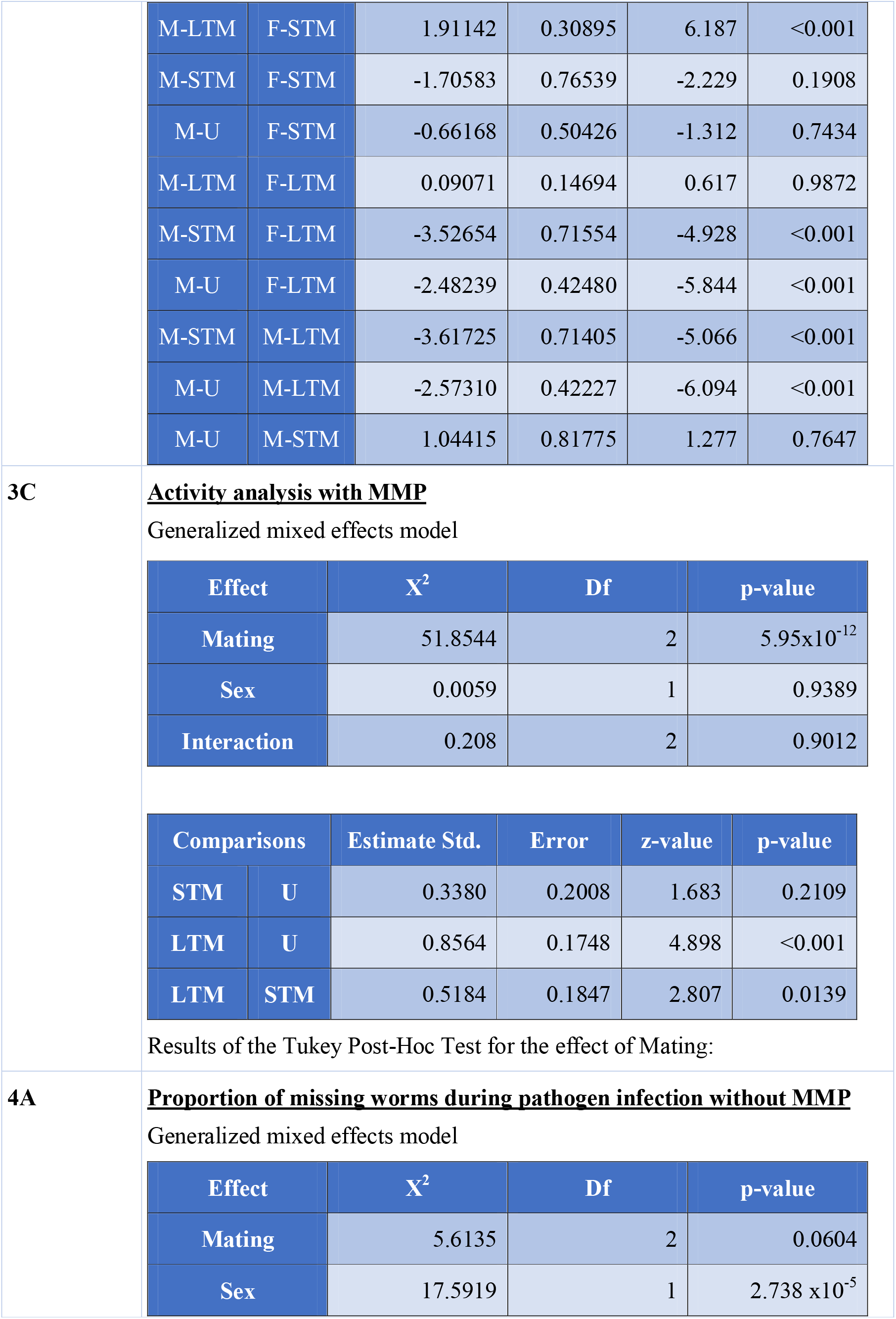
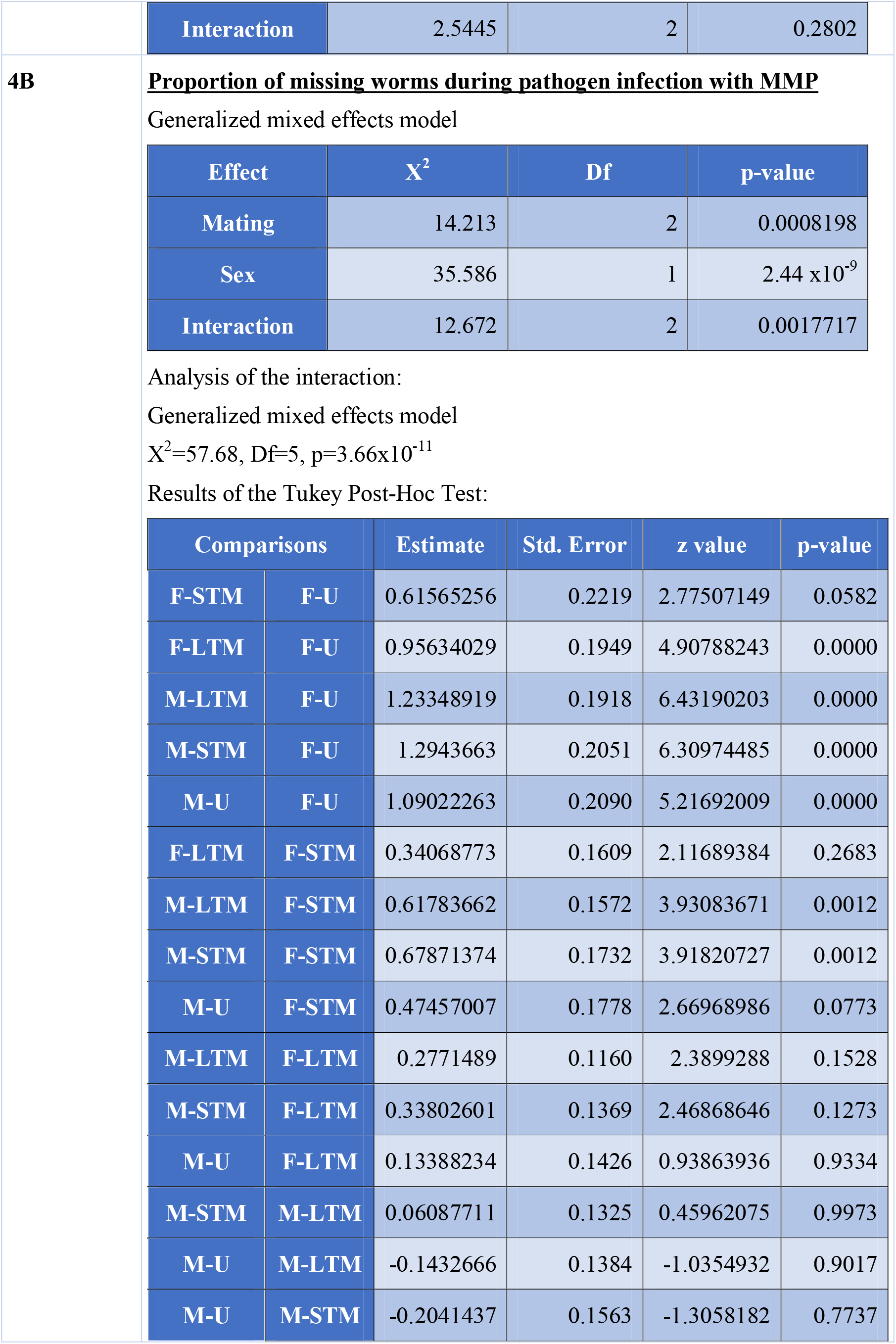
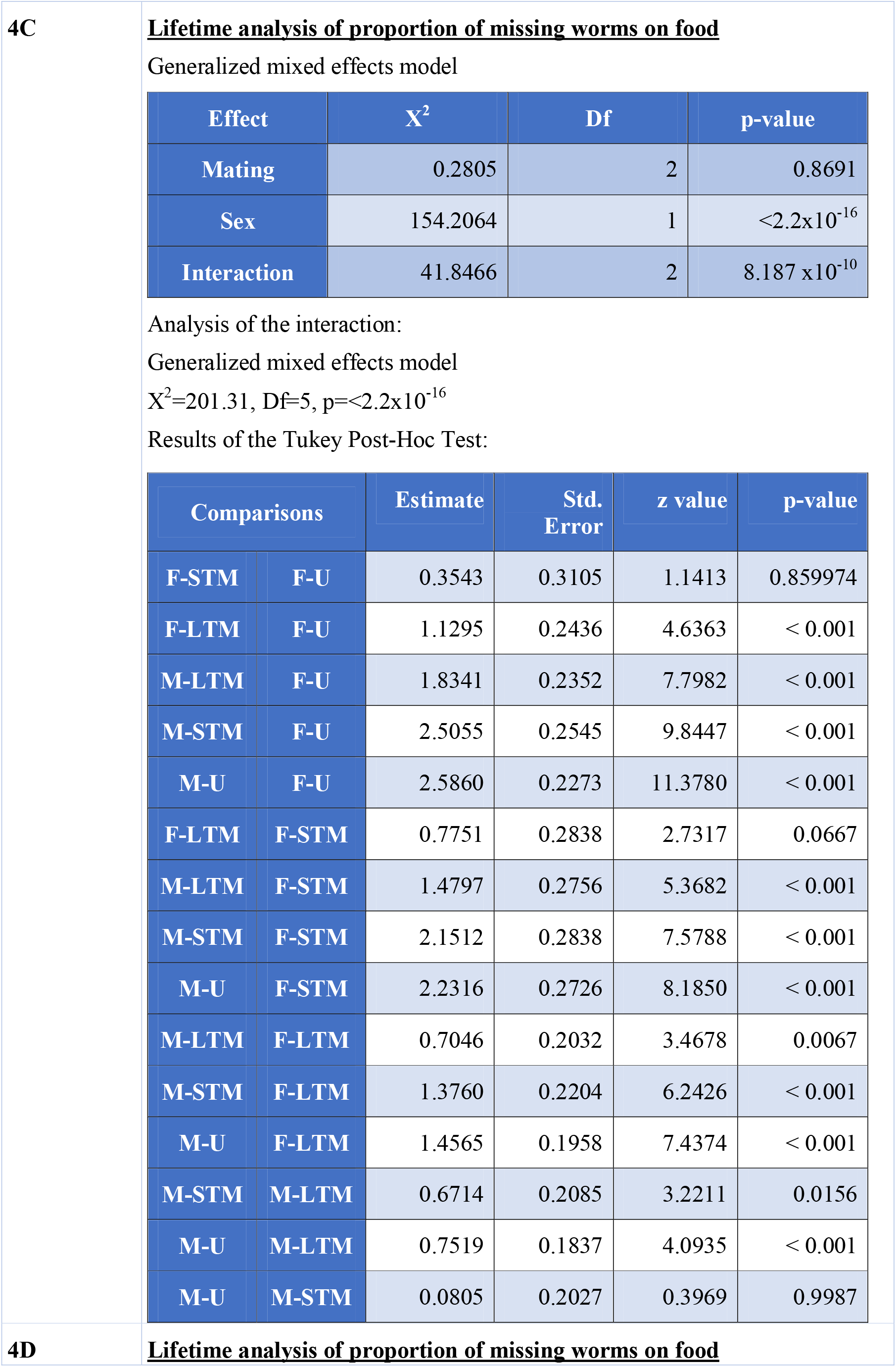
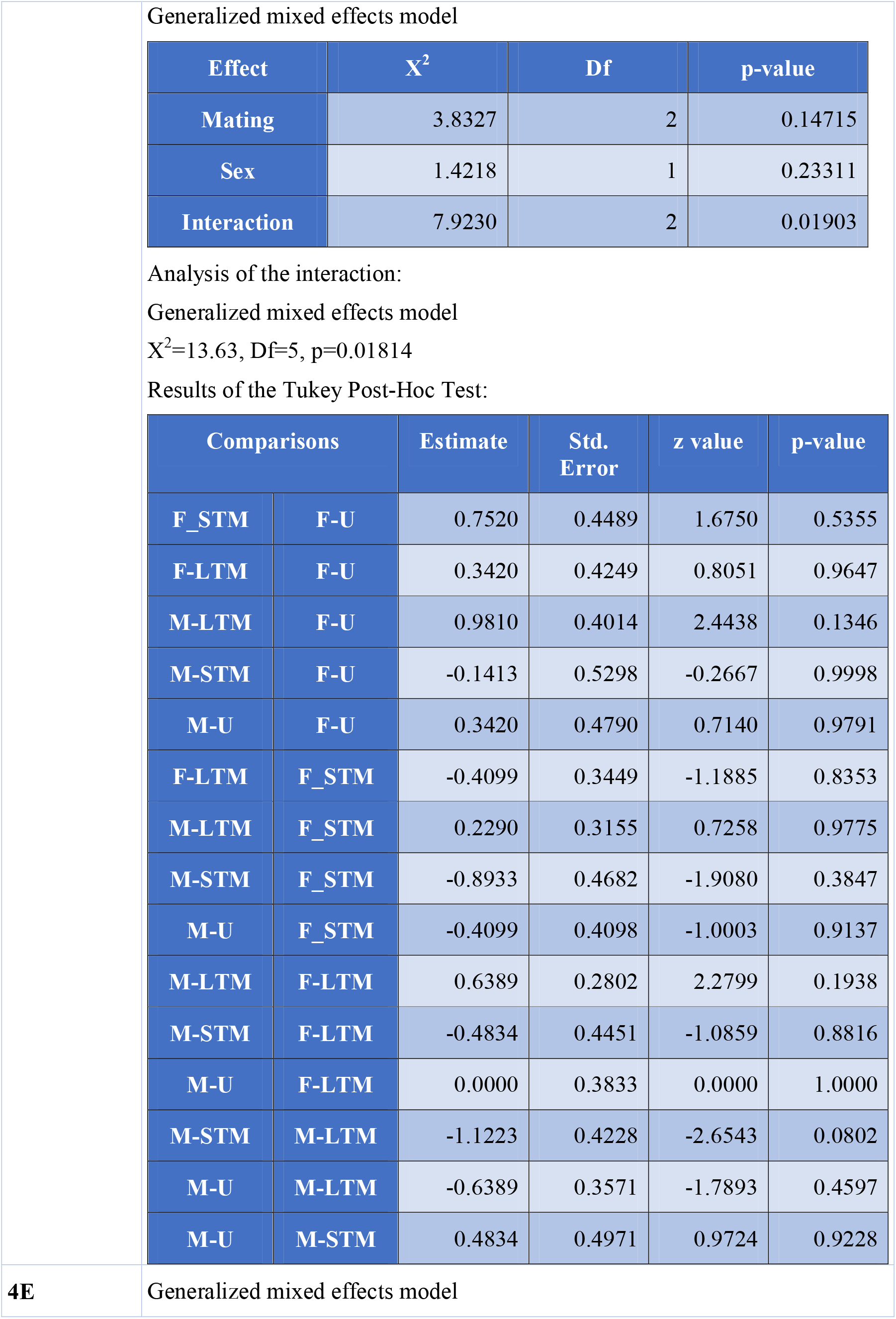
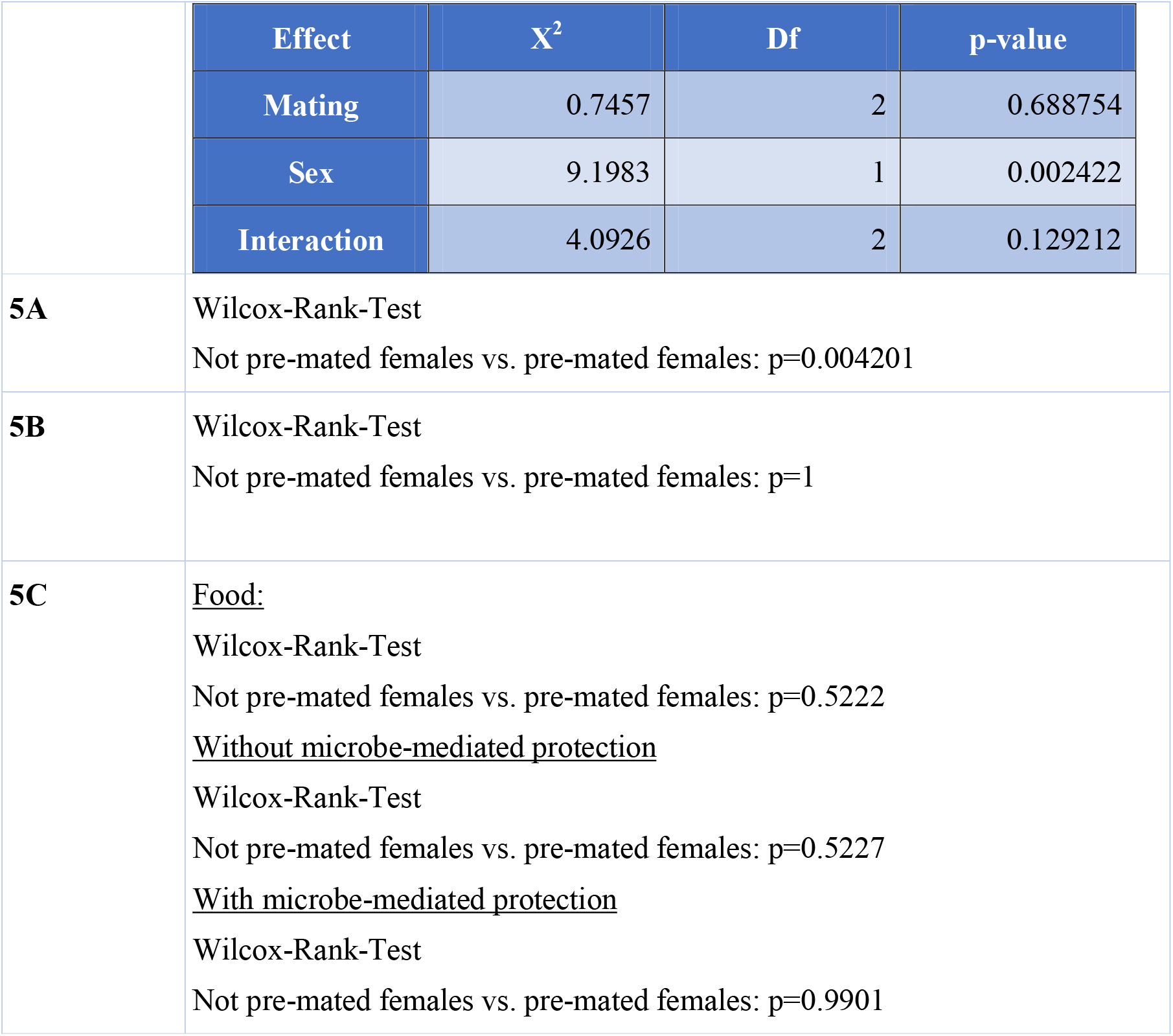
Summary of all statistical results. The following abbreviations were used: U=unmated, STM= short-term mated, LTM= Lifetime mated

